# Virus-like particle-delivered base editor collection to expand the genome engineering toolbox

**DOI:** 10.64898/2026.08.17.745336

**Authors:** Asfar Lathif Salaudeen, Trevor Shyiak, Carl G. de Boer

## Abstract

Virus-like particles (VLPs) enable transient, non-integrating delivery of CRISPR-Cas9 ribonucleoprotein cargo. Although VLPs have been reported for efficient DNA editing via base editors RNP delivery, the diversity of base editors tested as VLPs remains limited. We generated and benchmarked a panel of 12 base editors on the v5 eVLP backbone, targeting three genomic loci (HEK3, B2M, PDCD1) across five VLP dosages in LentiX-293T cells. Editing efficiency was generally dosage-dependent across all editors and varied by editor class and identity; PAM-flexible variants had lower editing efficiency than NGG-restricted counterparts, and the dual-function SPACE base editors showed reduced efficiency. We further characterized position-specific editing efficiencies and outcomes of the base editor VLP collection, revealing that a wide variety of mutation types are possible with the base editors in this collection.

## Background

The advent of genome engineering technologies over the last decade has revolutionized functional genomics. CRISPR-Cas9-based tools such as base editors and prime editors enable targeted, precise genome engineering^1^, facilitating disease modeling^2–4^, variant effect mapping^5,6^, and gene-editing-based therapeutics^7–10^. However, efficient delivery of these tools into cells remains a major bottleneck. Traditional CRISPR machinery delivery methods such as electroporation and lipid nanoparticles (LNPs) mediated deliveries enable transient transfection of plasmids, RNA or ribonucleoprotein (RNP) complexes, while lentiviral or transposon systems such as piggyBac mediate stable expression through integration^11–13^. However, these existing systems each carry their own limitations, including cargo constraints, inefficient cargo delivery, and permanent integration of mutagenic machinery into the genome^11,12^.

Virus-like particles (VLPs) have recently emerged as a delivery platform that combines the transduction efficiency of lentiviral systems with the transient activity of RNP-based delivery of CRISPR-Cas9 systems^14–19^ (Figure 1a). VLPs are produced by transfecting producer cells with lentiviral packaging components; however, unlike lentiviruses, VLP production does not use a transfer plasmid that integrates cargo into the recipient genome. Instead, a plasmid encoding Cas9 and a single guide RNA (sgRNA) enables the producer cell to assemble Cas9-sgRNA RNPs, which are then packaged into the particle itself (Figure 1a). Because VLPs retain the lentiviral envelope proteins that confer broad tropism, they allow efficient cargo delivery into otherwise hard-to-transfect cell types, while the RNP-based cargo preserves their transient, non-integrating nature post-transduction. Furthermore, as the CRISPR-Cas9 base editors and their sgRNA are typically packaged together as RNPs, multiple different base editors can be used simultaneously with minimal cross talk, making them well suited for orthogonal multiplexed editing at several genomic loci^20^.

**Figure 1:**
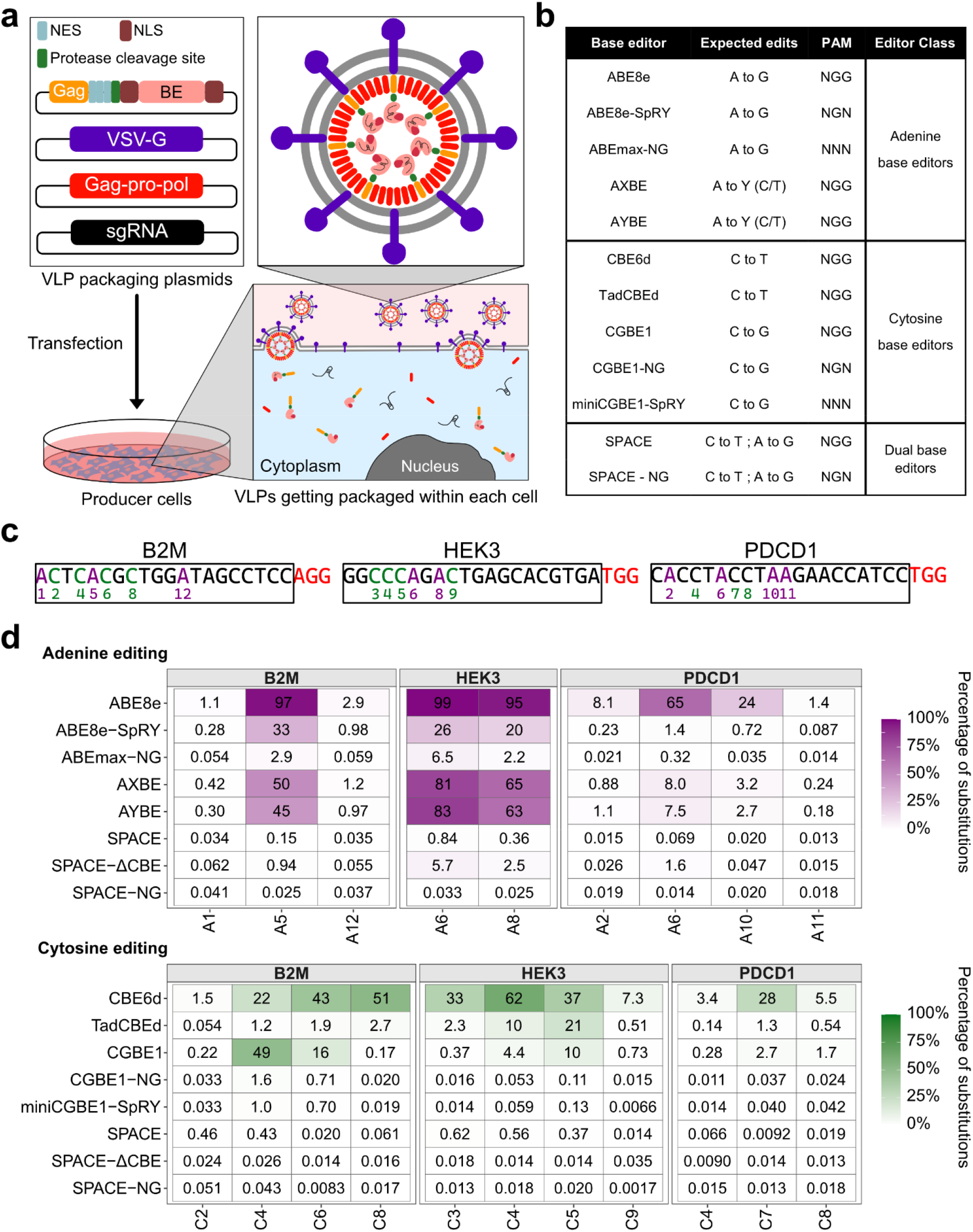
Base editor VLP production and editing efficiency of base editor VLP pane across three genomic loci. (a) Schematic of the engineered base editor virus-like particle (VLP) production. Producer cells are transfected with VLP packaging plasmids: Gag–propol, Gag–base editor (Gag-BE), VSV-G (envelope glycoprotein) and sgRNA expression plasmid. Producer cells then express sgRNAs and base editor proteins, which combine into RNPs. Base editors are fused to a Nuclear Export Signal (NES) to enable cytoplasmic localization and Gag for VLP packaging. Upon VLP infection of target cells, a protease cleavage site separates Gag and the NES from the base editor, and Nuclear Localization Signals (NLS) facilitate nuclear localization of the RNP. (b) Base editor panel used in this study, grouped by editing outcome and PAM compatibility. ABE8e VLP had been characterized in previous studies. (c) Target genomic sequences for the three loci assayed: B2M, HEK3, and PDCD1 with adenine (purple) and cytosine (green) positions relevant to editing numbered according to protospacer position. (d) Heatmaps of mean percentage of reads with substitutions for adenines and cytosines (top and bottom subplots) across the editable positions (x-axes) for B2 M, HEK3, and PDCD1 loci (left-right subplots), for all tested base editors (y-axes). Editing efficiency is calculated as the average of replicates (n=3) at a 10 μl VLP dose. Only positions with at least 1% editing in at least one of the base editors in that editing class are shown.

Despite this promise, VLP-mediated delivery has only been validated for a limited set of base editors, leaving most of the field’s expanding mutational and PAM-targeting toolbox untested and unexplored. Here, we generate and benchmark a collection of VLP-compatible base editors spanning diverse mutational spectra and PAM flexibilities, evaluating their editing efficiency and specificity across multiple genomic loci.

## Results and Discussion

To construct the base editor VLP collection, we assembled the backbone of the v5 eVLP from Raguram et al. 2025^19^ with 12 base editors with varying mutational spectra and PAM flexibility (Figure 1b). To generate the VLPs, we co-transfected viral packaging plasmids along with a VLP-compatible base editor plasmid and sgRNA expression plasmid for each target and base editor into the Lenti-X™ 293T cell line (Figure 1a), and then concentrated the VLPs from the media (Supplementary Figure 1, Methods). We targeted three loci within the human genome that had previously been used to benchmark base editing^21^: HEK3, B2M and PDCD1 (Figure 1c). We transduced the concentrated VLPs into Lenti-X™ 293T cells to test editing efficiency (Supplementary Figure 1, Methods).

We first confirmed that our three target loci were editable via base editor VLPs using the previously reported v5-ABE8e eVLP^16^, the best characterized editor in our panel, before extending testing to the remaining 11 base editors. We observed A-to-G editing at all three target loci, with efficiencies varying both between and within loci and some bases more prone to editing than others (Figure 1d, Supplementary Figure 2), consistent with the editing window previously reported for ABE8e^22^. We then tested the other 11 base editors in the same way (Adenine base editors: ABE8e-SpRY^23^, ABEmax-NG^24^, AYBE^25^, AXBE^26^; Cytosine base editors: CBE6d^27^, TadCBEd^21^, CGBE1^28^, CGBE1-NG^28^, and miniCGBE1-SpRY^28^; Dual (AC) base editors: SPACE^29^, and SPACE-NG^29^). As with ABE8e VLP, we observed varying efficiencies of base-position-specific editing both within and across the three loci (Figure 1d, Supplementary Figure 2). Editing activity varied by base editor even among those of the same class (e.g., CBE6d vs. TadCBEd; Figure 1d, Supplementary Figure 2), consistent with previous reports^27^. NGG-restricted base editors showed substantially higher editing efficiency than their PAM-flexible counterparts (ABE8e vs. ABE8e-SpRY; CGBE1 vs. CGBE1-NG vs. miniCGBE1-SpRY) (Figure 1d, Supplementary Figure 2), reinforcing that the expanded targeting range of PAM-flexible variants comes at an efficiency cost^30–32^. This trade-off is one that users will need to weigh against the need to target loci lacking canonical NGG sites. VLPs packaging dual base editors (SPACE) showed substantially lower editing efficiency than either adenine or cytosine base editors alone (Figure 1d, Supplementary Figure 2), potentially because the larger size of the base editor, with dual-deaminase proteins, reduces packaging or delivery efficiency into VLPs. To test this, we constructed an additional base editor, SPACE-ΔCBE, a truncated SPACE construct lacking the CBE (cytidine deaminase) component, leaving only the ABE domain, and observed a substantial increase in A-to-G editing across all three target loci relative to full-length SPACE (Figure 1d), supporting the hypothesis that base editor cargo size might limit VLP packaging. This suggests further cargo-specific VLP optimizations may be necessary for these base editors. Given the substantially reduced editing efficiency of the dual base editors across all three loci, we excluded all the dual base editors from subsequent analyses and focused on the 10 single-deaminase base editors.

**Figure 2:**
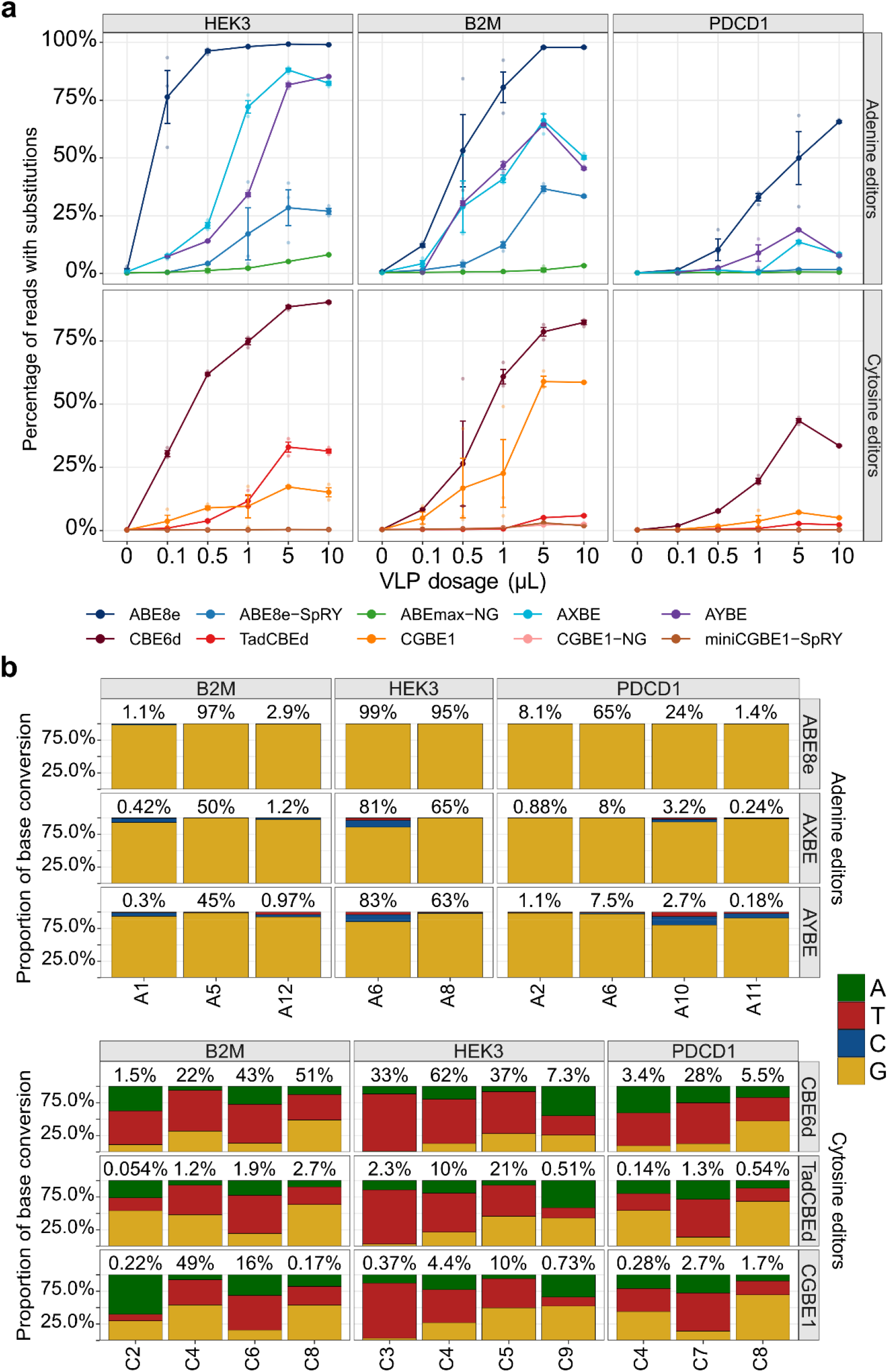
Dosage-dependent editing efficiency and base conversion outcomes of adenine and cytosine base editors delivered via VLPs. Percentage of reads with substitutions (y-axis) at target loci (columns) as a function of VLP dosage (x-axis), for adenine base editors (top row) and cytosine base editors (bottom row) across HEK3, B2M, and PDCD1 (left-right subplots). Solid points and error bars represent mean ± SD across replicates (n=3); colors denote individual base editors. (b) Base editing outcomes for the NGG base editors across the 3 target loci. Mean proportion of base conversion outcomes across replicates (n=3; y-axes) at edited positions (x-axes) for adenine base editors and cytosine base editors (top and bottom subplots) across B2M, HEK3, and PDCD1 (left-right subplots), at the highest VLP dosage (10 μl). The base positions shown (x-axes) were observed to have at least 1% editing within one of the base editors in that editing class.

To determine how editing efficiency scales with VLP input, we tested each base editor across a five-point dosage range (0.1, 0.5, 1, 5 and 10 μl) at all three loci (Supplementary Figure 1). In addition to differences between editors and the edited loci, editing efficiency for a given base editor was itself highly dependent on VLP dosage. We observed increasing editing efficiency with increasing VLP input for both adenine and cytosine editors. However, the magnitude and shape of this dose-response relationship varied by editor and locus (Figure 2a, Supplementary Figure 3). Given the dosage-dependent relationship between VLP concentration and editing efficiency observed here, we recommend that users titrate VLP dose empirically for their locus of interest and apply the minimum concentration that yields appreciable editing, to decrease the risk of off-target activity^33,34^.

We next looked at the substitution types produced by each editor. ABE8e made only the classical transition edits of A-to-G as expected for this editor (Figure 2b). The same was true for the PAM-flexible ABEs (ABEmax-NG and ABE8e-SpRY) on bases with appreciable (≥1%) edits (Supplementary Figure 4). Both AYBE and AXBE, designed to produce A-to-C/T edits, predominantly produced A-to-G edits, but resulted in A-to-C/T edits occasionally as well (Figure 2b). CGBE1, meanwhile, designed to favour C-to-G edits, produced a fairly even distribution of the three alternative bases, although this varied substantially by position. In addition to CGBE1, CBE6d and TadCBEd VLPs also produced a variety of mutation outcomes despite neither the CBE6d^27^ nor TadCBEd^21^ source publications reporting C-to-G/A editing. However, related TadA-derived CBE variants (CBE-T1.14, CBE-T1.46, CBE-T1.52) show C-to-non-T editing in the range of ∼1–15% depending on target site^35^. Of the total amount of edits, we observed a proportion of C-to-non-T edits ranging from 13–70% for CBE6d and 18–73% for TadCBEd, across the three target loci and their edited bases (bases with ≥1% overall editing on at least one of the base editors). (Figure 2b, Supplementary Table 1). A comparable discrepancy in CBE editing purity has been reported for VLP-packaged AncBE4max (Addgene #112094) by a previous study^20^. Although AncBE4max carries a C-terminal 2×UGI cassette, which normally suppresses base excision repair of the uracil intermediate and thereby enforces high C-to-T purity, VLP packaging of this editor led to a marked shift toward C-to-non-T outcomes. This effect was reversible: co-delivering supplemental UGI to recipient cells via a separate VLP restored C-to-T editing^20^. We propose two non-exclusive explanations for this loss of purity: the amount of UGI incorporated per VLP particle is insufficient to fully saturate and inhibit endogenous Uracil-DNA glycosylase activity, leaving more uracil intermediates available for C-to-non-T editing outcomes; or the UGI domain is proteolytically cleaved from the base editor during eVLP maturation, when the viral protease processes the Gag-Pol-cargo fusion, leaving the editor UGI-deficient. Another alternative is the site dependence of editing outcome, which is clear in our data, and seems to be true for all tested base editors capable of generating multiple substitution types. This did not appear to be specific to protospacer position, as even within the same position (e.g., A6 for HEK3 and PDCD1, C4 for all three loci) both editing efficiency and mutation outcome distribution differed substantially (Figure 2b). Notably, these two positions differ in local trinucleotide context (CAG and TAC for A6 and CAC, CCC, and CCT for C4), suggesting that local sequence context, in addition to window position, modulates the mutation identity. This position-specific base editing outcome pattern is reminiscent of the context-dependent mutability described in cancer mutational signatures^36^. These results further reinforce that the context and position of an editable base influence not only the overall editing efficiency but also the substitution type.

In addition to base editing activity, several base editor VLPs also produced deletions within the editing window (Supplementary Figure 5). The deletion frequency closely followed the editing frequency (Supplementary Figure 6), consistent with increased indels accompanying high base editing events^37^. However, the ratio of substitutions to deletions differed substantially between base editors (Supplementary Figure 7), with CBEs in particular having a propensity to result in deletions.

## Conclusion

Base editor VLPs address key limitations of traditional gene editing delivery methods by enabling transient, tunable, and efficient base editing activity. Here, we expanded the diversity of the VLP-compatible base editor toolkit by packaging 12 different base editors spanning a range of mutational spectra and PAM requirements, and characterized their dosage-dependent editing activity, including their mutational spectra, across three genomic loci. We observed that, for certain base editors, the editing outcomes varied not only by position within the editing window but also, we infer, by local sequence context. Hence, users should also empirically profile editing outcomes, not just efficiency, at their locus of interest rather than assuming a fixed outcome across targets. Given the relative ease of packaging target-specific sgRNAs and base editors into VLPs, this platform could support scalable multiplexed genome engineering, including simultaneous packaging and/or delivery of multiple sgRNAs and even base editors. VLP-delivered CRISPR-Cas9 editors can also be further engineered for cell-type specificity^16^, to edit harder-to-transduce cell types, including human hematopoietic stem cells and other primary cells^15^, and for in vivo delivery^14,17^, broadening their potential as a gene therapy delivery platform.

## Methods

### Assembling the base editor VLP collection constructs

The base editor VLP collection was generated via Gibson assembly of the base editor inserts amplified from the corresponding base editor expression plasmid and assembled into the VLP backbone amplified from v5-eVLP-ABE construct from Raguram et al, 2025^19^ (Addgene: 228467). The assembly mix was then transformed into NEB Stable cells via heat-shock transformation according to manufacturer’s protocol. Briefly, 50 μl of competent cells were transformed with 2 μl of the assembled DNA followed by incubation on ice for 5 mins. Then the cells were heat-shocked for 60 sec at 42 °C and then placed on ice for 5 mins. The cells were then diluted with LB media and shaken at 30 °C for 1 hour for recovery. After recovery growth, transformed cells were then plated on LB agar with 100 µg/mL ampicillin and incubated at 30 °C for 16 hours. Colonies were picked the next day, cultured overnight, miniprepped, and the plasmid sequence was verified using Oxford Nanopore Technology sequencing via Plasmidsaurus. All the primers used for amplifying the base editor inserts and VLP backbone are attached in the Supplementary Materials.

### sgRNA cloning

Single guide RNAs (sgRNAs) targeting HEK3, B2M, and PDCD1 were cloned using a Polymerase Cycling Assembly (PCA)-based Golden Gate strategy adapted from RAPID-DASH protocol^38^. U6 promoter and scaffold/terminator fragments were PCR-amplified from a U6-sgRNA vector (MLM3636) using Phusion High-Fidelity polymerase (Thermo F531) with manufacturer’s protocol. For each locus, a 59-nt spacer oligonucleotide encoding the 20-nt protospacer (HEK3: GGCCCAGACTGAGCACGTGA; B2M: ACTCACGCTGGATAGCCTCC; PDCD1: CACCTACCTAAGAACCATCC) flanked by homology arms to the promoter and scaffold fragments was assembled with the U6 promoter and terminator/scaffold fragments by PCA, generating sgRNA units with flanking type IIS (BsaI) restriction sites. The PCA reactions were set up as follows: 1 µL U6 promoter (5 ng), 1 µL gRNA terminator scaffold (5 ng), 2 µL spacer oligo (1 µM), 1.25 µL forward primers (10 µM), 1.25 µL reverse primers (10 µM), 25 µL Phusion High-Fidelity PCR Master Mix (Thermo F531), and nuclease-free water to a total volume of 50 µL. The following setting was used in thermocycler: 98 °C for 3 min, 35 cycles of 98 °C for 10 s, 52 °C for 30 s, and 72 °C for 12 s which was followed by a final 72 °C for 8 min. PCA products were confirmed by agarose gel electrophoresis and purified with AMPure XP beads (1×). Purified sgRNA units were cloned into the Golden Gate destination vector pAL11 in a one-pot Golden Gate reaction as follows: 100 ng of the destination vector, 500 ng of the purified sgRNA units, 1 µL of T4 DNA ligase (400 U/µL, NEB M0202S), 1.5 μl FastDigest Eco31I (Thermo FD0293) (BsaI isoschizomer), 2 µL of 10X T4 DNA ligase buffer (NEB), and nuclease-free water up to 20 µL. The following setting was used in the thermocycler: 5 min at 37 °C and 5 min at 16 for 30 cycles followed by 37 °C for 10 min. The restriction enzyme was then inactivated at 75 °C for 10 min. 2 µL of the assembly mix was then transformed into NEB Stable cells via heat-shock transformation according to manufacturer’s protocol. Transformed cells were then plated on LB agar with 100 µg/mL ampicillin and incubated at 30 °C for 16 hours. Colonies were picked the next day, cultured overnight, miniprepped, and the plasmid sequence was verified using Oxford Nanopore Technology sequencing via Plasmidsaurus. All the primers used for assembling sgRNA constructs are attached in the Supplementary Materials.

### Cell culture conditions

HEK Lenti-X™ 293T (Takara 632180) were cultured in DMEM, high glucose (Gibco 11965092) supplemented with 10% fetal bovine serum and 1% penicillin and streptomycin as growth medium. Cells were cultured in 37 °C with 5% carbon dioxide.

### Base editor VLP generation and concentration

BE-VLPs were produced by transiently transfecting Lenti-X™ 293T cells with the corresponding packaging plasmids (Supplementary Figure 1a). Lenti-X™ 293T cells were seeded in 6-well plates at a density of 800,000 cells per well. Each well produced a unique BE-VLP with one of 13 base editors (ABE8e, ABE8e-SpRY, ABEmax-NG, AYBE, AXBE, CBE6d, CGBE1, CGBE1-NG, miniCGBE1-SpRY, TadCBEd, SPACE, SPACE-ΔCBE, and SPACE-NG) targeting one of 3 loci (B2M, HEK3, and PDCD1). 12–16 hours after seeding, growth medium was replaced with 3 ml fresh growth medium. 12 hours later, cells were transfected with 3,109 ng of plasmid DNA at a 1:8.4:2.8:11 mass ratio of pMD2.G (vesicular stomatitis virus G (VSV-G)) (Addgene: 12259), eVLP-v5-Gag-Pro-Pol (Addgene: 228466), one of the base editor VLP plasmids, and an sgRNA expression plasmid (134, 1,126, 375, and 1,474 ng, respectively). Plasmids were diluted in a solution comprised of 300 μl OptiMEM (Gibco 31985062) and 12.4 μl Transporter5 transfection reagent (Kyfora Bio 26008) (1 μg DNA: 4 μL transfection reagent ratio), vortexed briefly, and incubated at room temperature for 20 minutes to allow transfection complex formation before addition to the cells. Transfection medium was replaced with 3 ml fresh growth medium 24 hours post-transfection. BE-VLP-containing supernatant was collected 48 hours after this media change, isolated from cells and debris by centrifugation (500 × g, 5 min), and filtered through a 0.45 μm PVDF syringe filter (Millipore SLHVR33RS). Filtered supernatant was concentrated by adding 1 ml of 4X Lenti Concentrator (80% (w/v) polyethylene glycol 8000 (PEG-8000) (Fisher BP233100) and 14% (w/v) NaCl suspended in 100 ml of 1×PBS, pH 7.4) and incubating overnight at 4 °C with continuous end-over-end rotation, followed by centrifugation at 1600 × g for 60 minutes at 4 °C. The supernatant was carefully removed, and the pelleted BE-VLPs were resuspended in PBS at 100X concentration relative to the original culture volume. Concentrated BE-VLPs were aliquoted and stored at −80 °C.

### Base editor VLP transduction and gDNA isolation

Lenti-X™ 293T cells were plated in 96-well plates at a density of 5000 cells per well for each of the base editor x dosage condition across three target loci in triplicates (Supplementary Figure 1b). 12–16 hours after seeding, growth medium was replaced with 3 ml fresh growth medium containing final concentration of polybrene (Sigma Aldrich TR-1003-50) to aid in VLP transduction. Each of the 39 BE-VLPs, with a unique base editor and target locus, were transduced directly to the culture media in each well at six volumes (0, 0.1, 0.5, 1, 5, 10 μl) in triplicate. 24 hours after transduction, growth medium was replaced with 3 ml fresh growth medium. 48 hours after the media change, gDNA was isolated from the cells as follows: cells were washed once with PBS, and lysed in 60 μL of lysis buffer (10 mM Tris-HCl pH 8.0, 0.05% sodium dodecyl sulfate (SDS), and 25 μg mL-1 Proteinase K (Thermo EO0491)) at 37 °C for 1 h, followed by heat inactivation at 80 °C for 30 min.

### Target amplification and high-throughput sequencing

For each sample, the gDNA-containing cell lysate was used to amplify the respective target loci in two steps to prepare the sequencing library (Supplementary Figure 8). Primers were designed to both amplify the three target loci (B2M, HEK3, and PDCD1) and additionally barcode each target by replicate to enable pooling across replicates. This was accomplished by designing three reverse primers for each target, and nine forward primers, each primer amplifying a target, and adding a replicate-specific barcode. PCR1 reactions for each combination of base editor, target, dosage and replicate were set up as follows: 1 μl cell lysate, 0.25 μl 10 μM replicate and target-specific forward primers (0.25 μM), 0.25 μl 10 μM target-specific reverse primers (0.25 μM), 0.5 μl 4X SYBR™ Green I (Thermo S7563) (0.2X), 5 μl Platinum SuperFi II polymerase (Thermo 12369010), and nuclease-free water to a total volume of 10 μl. Thermocycler conditions for PCR1: 95 °C for 3 min; 24 cycles of 95 °C for 10 s, 60 °C for 10 s, and 72 °C for 15s; 72 °C for 5 min. The cycle number was determined by qPCR, stopping amplification when the libraries were detectably in exponential amplification. This yielded amplicons for each base editor, target, and dosage, barcoded by replicate. After PCR1, the corresponding replicates were pooled together for every unique (target × base editor × dosage) combination, which were then distinguished with Illumina indices. Each pooled library was used as an input for PCR2, with forward and reverse primers containing i5 and i7 index barcodes for paired-end dual-index Illumina sequencing that were unique for each (base editor × dosage) combination. PCR2 reactions were set up as follows: 0.5 μl pooled PCR1 product, 0.5 μl 10 μM i5-indexed forward primers (0.5 μM), 0.5 μl 10 μM i7-indexed reverse primers (0.5 μM), 0.5 μl 4X SYBR™ Green I (Thermo S7563) (0.2X), 5 μl NEBNext® Ultra™ II Q5® Master Mix (NEB M0544), and nuclease-free water to a total volume of 10 μl. Primers for PCR2 containing the i5 and i7 Illumina barcodes (unique dual index or UDI) were generated using the tool GIL^39^. Thermocycler conditions for PCR2: 98 °C for 3 min; 5 cycles of 98 °C for 10 s and 65 °C for 75 s; 65 °C for 5 min. qPCR was again used to determine when to stop amplification. After PCR2, amplicons were pooled and gel purified in a 1% agarose gel using the GeneJET Gel Extraction Kit (Thermo K0691). The final sequencing library was thus distinguished by the UDIs indicating the base editor and dosage, an internal barcode representing the replicate, and the loci distinguished by their unique sequences. Sample concentration was quantified using the Qubit High-Sensitivity Assay Kit (Thermo Q33231). Samples were then sequenced on an Illumina NovaSeq X Plus (PE150), obtaining a median of 570803 reads per UDI (each comprised of 9 locus-replicate combinations) (Supplementary Figure 9a). UDIs corresponding to the 0 μL dosages of AYBE and SPACE-ΔCBE returned no reads, and so were excluded from analysis (although we note that since no VLP was administered in these, they are effectively identical to the other 0 μL samples for other base editors). All the primers and replicate-barcode sequences used here are listed in the Supplementary Materials.

### High-throughput sequencing data analysis

Paired-end reads were demultiplexed based off of the UDI into their respective sample condition (base editor × dosage), with all target loci and biological replicates pooled together. Paired-end reads were merged into single amplicon-spanning reads using PEAR (v0.9.6)^40^ with default parameters. Read quality was assessed with FastQC (v0.12.1)^41^ both before and after merging, and reports were aggregated with MultiQC (v1.33)^42^, verifying the samples were all of sufficient quality. Merged reads were demultiplexed into per-replicate FASTQ files using cutadapt (v5.2)^43^, matching a 5′-anchored replicate-specific barcode (error rate = 0.17, no indels permitted). Upon replicate demultiplexing, there were 172,366 reads per sample condition replicate, on average (Supplementary Figure 9b). Base editing outcomes were quantified for each sample condition (base editor + dosage)–replicate combination using CRISPRessoPooled (CRISPResso2 (v2.3.4)^44^) against a multi-amplicon reference file, with base-editor output mode and the following parameters adjusted: --plot_window_size: 10, --quantification_window_center: -10 and --quantification_window_size: 10. Target level demultiplexing via CRISPRessoPooled yielded a median of 34,672 reads for HEK3, 54,972 reads for B2M and 72,824 reads for PDCD1 (Supplementary Figure 9c). To compare editing outcomes across dosages within each base editor, replicate-matched CRISPRessoPooled outputs for all doses of a given base editor were aggregated using CRISPRessoAggregate. Base editor efficiencies are reported as the percentage of sequencing reads containing a certain base conversion within the protospacer sequence (quantification window). All the plots and statistical analysis were done in R 4.4.3 using tidyverse (v2.0.0).

### Ethics approval and consent to participate

Not applicable. This study did not involve human participants, human data, human tissue, or animals; all experiments were performed in the HEK293T (Lenti-X™ 293T) immortalized cell line.

## Consent for publication

Not applicable.

## Availability of data and materials

Raw sequencing reads can be accessed at SRA (PRJNA1512594); All the plasmids generated in this study can be accessed in Addgene. All the primers used in this study can be accessed in the Supplementary Materials. All the codes used to process sequencing data and generate plots are available at GitHub (https://github.com/de-Boer-Lab/VLP_BEs/).

## Authors’ contributions

The project idea was conceived by A.L.S. and C.G.D. A.L.S. generated the base editor VLP constructs. A.L.S. and T.S. performed the VLP generation and transduction experiments. A.L.S. performed the sequencing read processing and Crispresso2 analysis. A.L.S. and T.S. wrote the code for data plotting. A.L.S., T.S., and C.G.D. wrote the manuscript.

## Acknowledgements

v5 ABE-eVLP (Addgene plasmid #228467; http://n2t.net/addgene:228467; RRID:Addgene_228467), v5 eVLP gag-pro-pol (Addgene plasmid #228466; http://n2t.net/addgene:228466; RRID:Addgene_228466), ABE8e (Addgene plasmid #138489; http://n2t.net/addgene:138489; RRID:Addgene_138489), NG-ABEmax (Addgene plasmid #124163; http://n2t.net/addgene:124163; RRID:Addgene_124163), pCMV-SpRY-ABE8e (Addgene plasmid #185671; http://n2t.net/addgene:185671; RRID:Addgene_185671), SpCas9-CBE6d-V106W (Addgene plasmid #215827; http://n2t.net/addgene:215827; RRID:Addgene_215827), and p2T-cmv-SpCas9-TadCBEd-V106W-BlastR (Addgene plasmid #193849; http://n2t.net/addgene:193849; RRID:Addgene_193849) were a gift from David Liu. AXBEv2 (Addgene plasmid #204606; http://n2t.net/addgene:204606; RRID:Addgene_204606) was a gift from Dali Li. AYBEv3 (Addgene plasmid #193967; http://n2t.net/addgene:193967; RRID:Addgene_193967) was a gift from Huawei Tong. CGBE1 (pRZ3885) (Addgene plasmid #140252; http://n2t.net/addgene:140252; RRID:Addgene_140252), CGBE1-NG (pBM1246) (Addgene plasmid #140256; http://n2t.net/addgene:140256; RRID:Addgene_140256), miniCGBE1-SpRY (pBM1571) (Addgene plasmid #170105; http://n2t.net/addgene:170105; RRID:Addgene_170105), SPACE (pRZ1813) (Addgene plasmid #140242; http://n2t.net/addgene:140242; RRID:Addgene_140242), and SPACE-NG (pRZ5150) (Addgene plasmid #140244; http://n2t.net/addgene:140244; RRID:Addgene_140244) were a gift from Keith Joung.

