## Supplementary figures and table for "Virus-like particle-delivered base editor collection to expand the genome engineering toolbox"

### **Supplementary Material**

1. Supplementary Figure 1
2. Supplementary Figure 2
3. Supplementary Figure 3
4. Supplementary Figure 4
5. Supplementary Figure 5
6. Supplementary Figure 6
7. Supplementary Figure 7
8. Supplementary Figure 8
9. Supplementary Figure 9
10. Supplementary Table 1

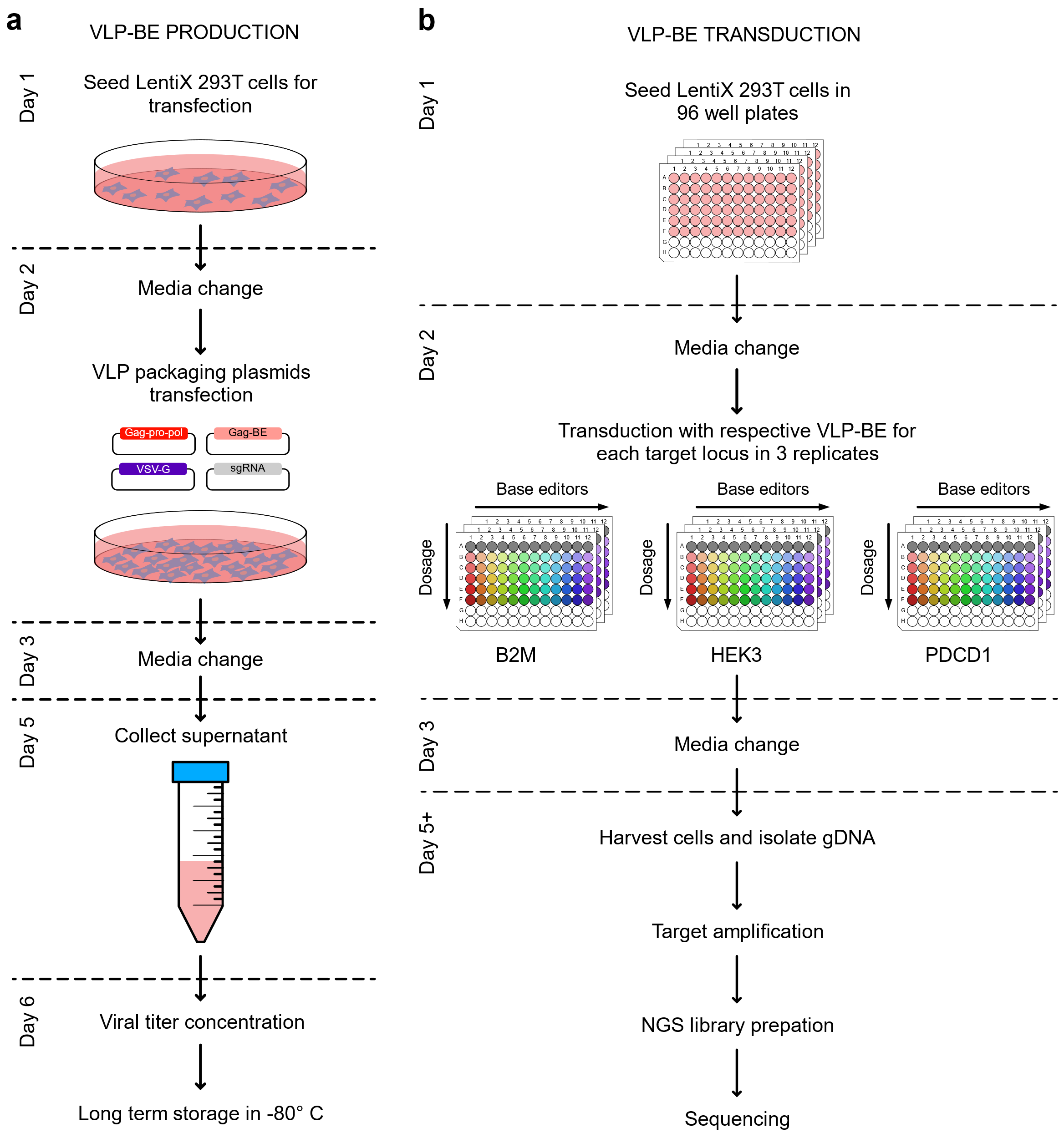

**Supplementary figure 1: VLP-BE production and transduction workflow.** (a) VLP-BE production timeline. LentiX 293T cells are seeded (Day 1) and co-transfected with the four VLP packaging plasmids: Gag–pro-pol, Gag–BE (base editor fusion), VSV-G envelope, and sgRNA expression plasmid (Day 2). Media is changed (Day 3), VLP-containing supernatant is collected (Day 5), and VLPs are concentrated and stored at −80 °C (Day 6). (b) VLP-BE transduction workflow. LentiX 293T cells are seeded in 96-well plates and transduced in triplicate with each base editor's VLP across the five-point dosage series, separately for the B2M, HEK3, and PDCD1 target loci. Following media change, genomic DNA is harvested, target loci are amplified, NGS libraries are prepared, and amplicons are sequenced. This panel corresponds to the workflow underlying every editing measurement reported.

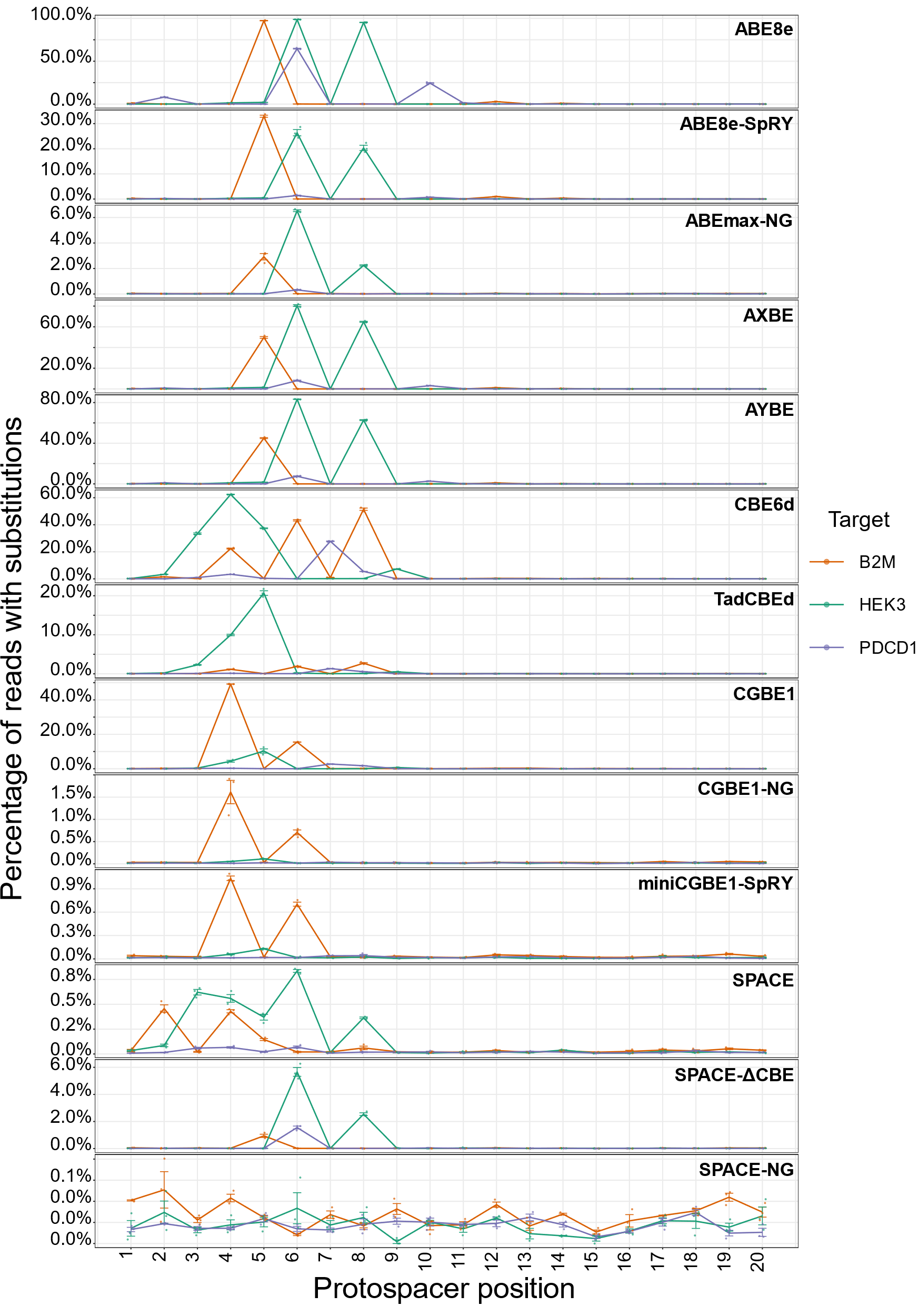

**Supplementary Figure 2: Base editor activity across the protospacer for each base editor and target locus.** Percentage of sequencing reads carrying a base substitution (y-axes) at each protospacer position (x-axes) is plotted for each of the 13 base editors in the panel showcasing their editing windows for the three target loci: HEK3, B2M, PDCD1 (colors). Points show the mean percentage of substituted reads (y-axes), with error bars representing the SEM across replicates (n=3).

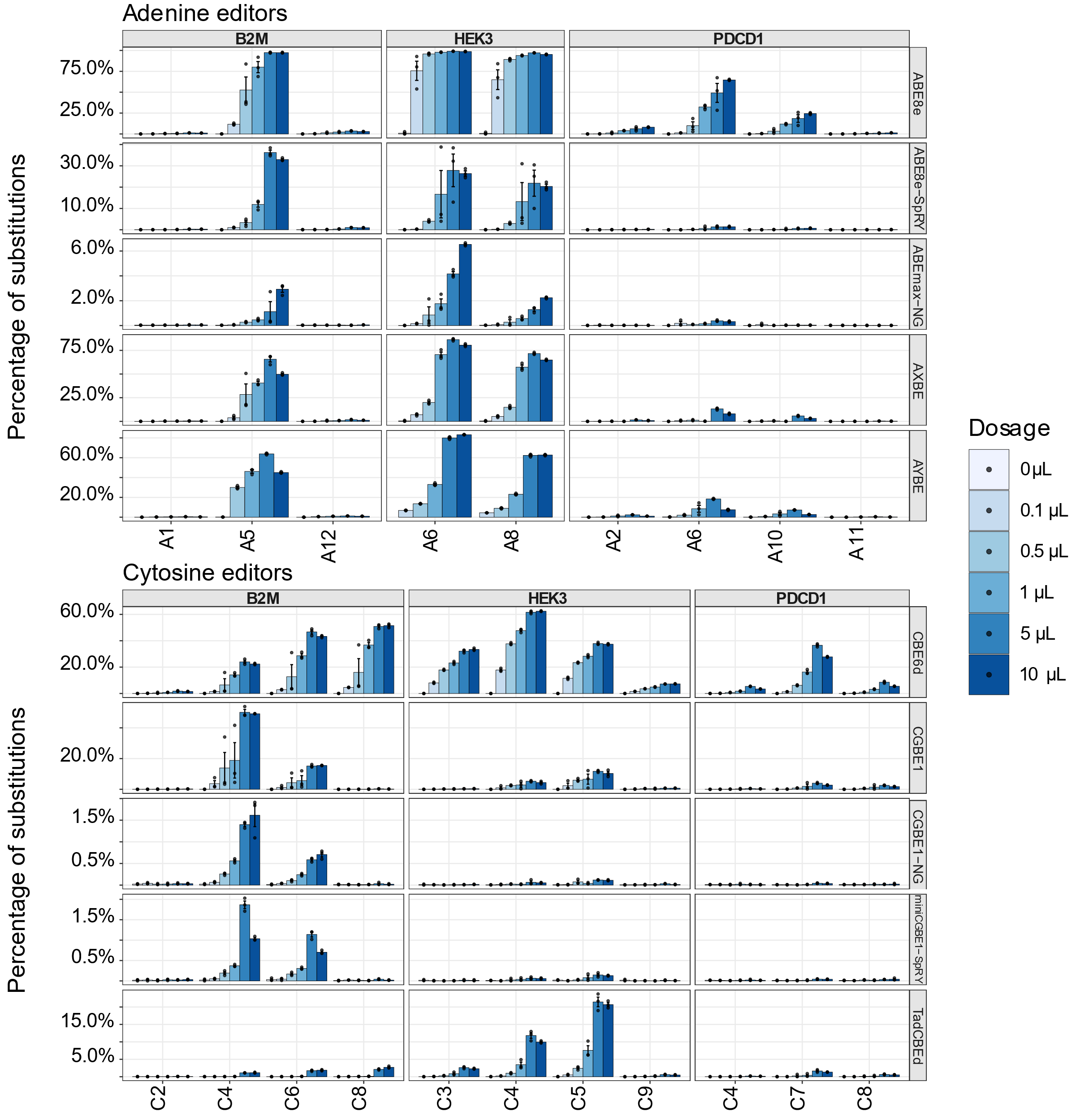

**Supplementary Figure 3: Dosage-dependent base-editing efficiency at individual adenine and cytosine positions across target loci.** Percentage of sequencing reads with substitutions (y-axes) is shown for each editable adenine (A) or cytosine (C) position within the protospacer (x-axes) at each of the three target loci: B2M, HEK3, and PDCD1 (left – right subplots) for each base editors within the adenine and cytosine base editor classes (top – bottom subplots). Bars represent mean editing percentage (y-axes) at each of six VLP dosages, with error bars representing SEM across replicates (n=3). The base positions shown (x-axes) were observed to have at least 1% editing within the base editors in that editing class. AYBE and SPACE-ΔCBE do not have a 0 µL data point.

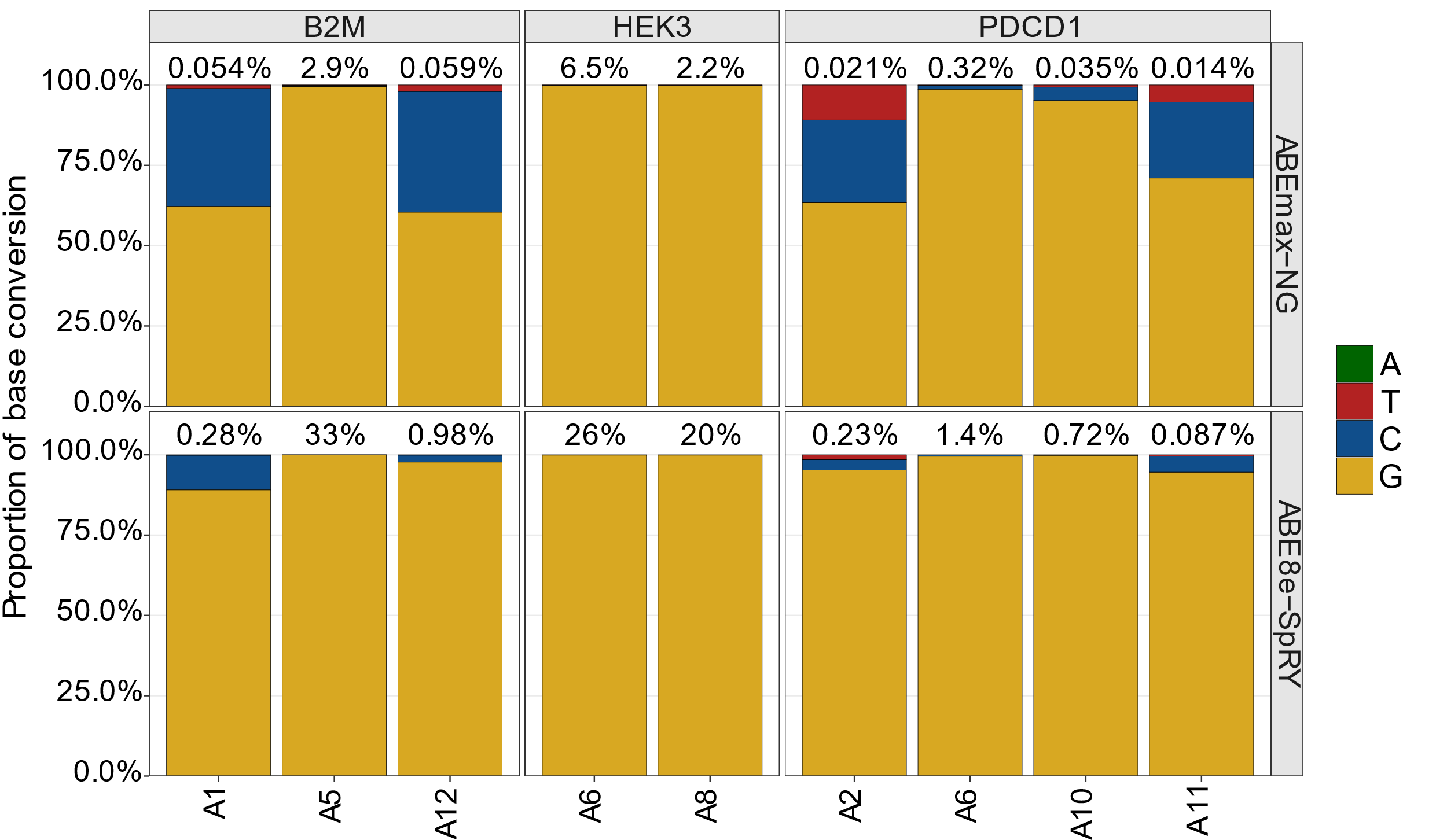
**Supplementary Figure 4: Base editing outcomes for the PAM-flexible ABE base editors across the 3 target loci.** Mean proportion of base conversion outcomes across the replicates (n=3; y-axes) at edited positions (x-axes) for PAM-flexible adenine base editors (top- bottom subplots) across the three target loci: B2M, HEK3, and PDCD1 (left – right subplots), at the highest VLP dosage (10 μl). The base positions shown were observed to have at least 1% editing within one of the base editors in ABE editing class.

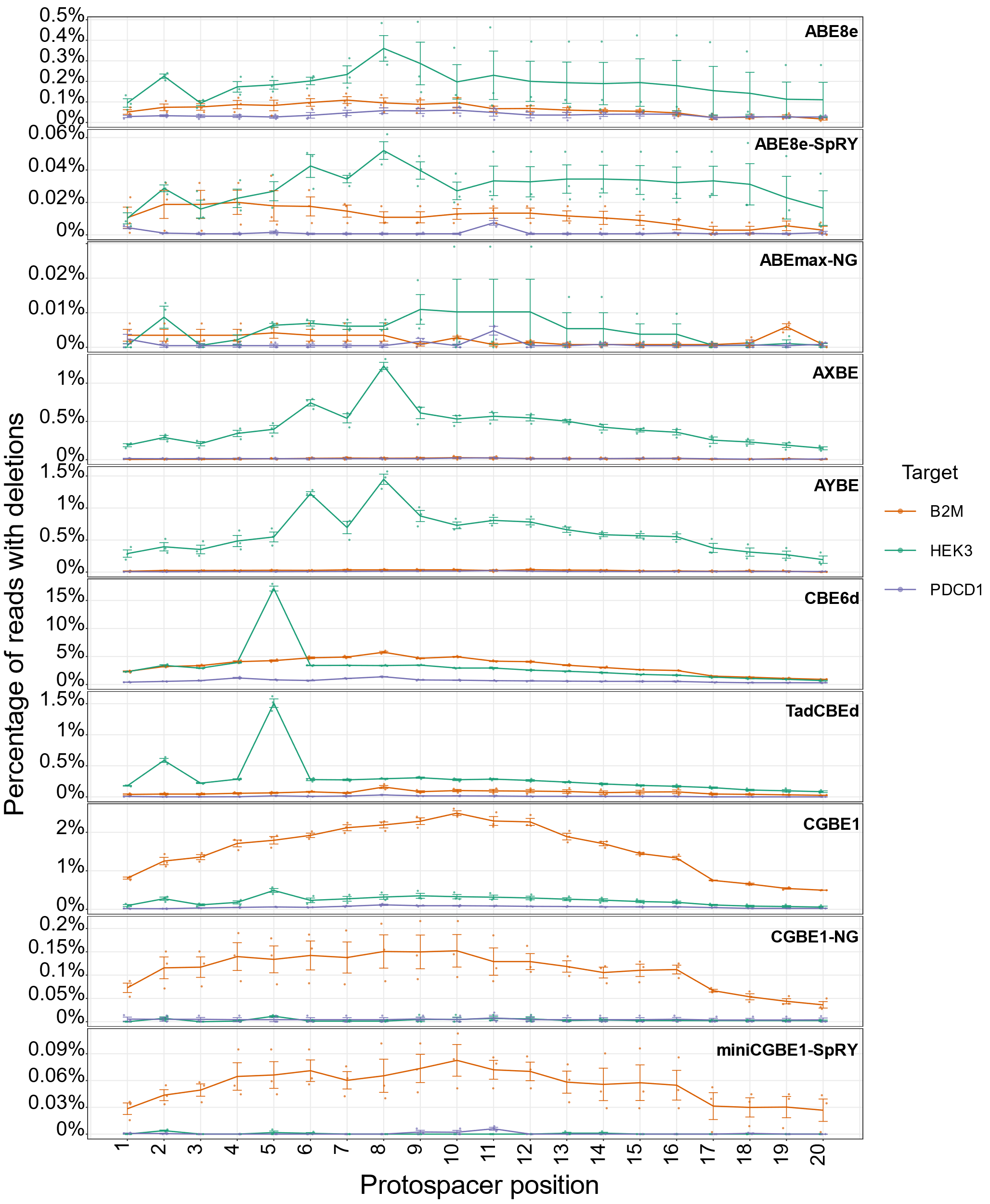

**Supplementary Figure 5: Deletion frequency across the protospacer for each base editor and target locus.** Percentage of sequencing reads containing a deletion (y-axes) overlapping each protospacer position (x-axes) is plotted for 10 base editors in the panel (rows; dual base editors not shown here), for the three target loci: B2M, HEK3, and PDCD1 (colors). Points show mean deletion percentage (y-axes), with error bars representing SEM across replicates (n=3).

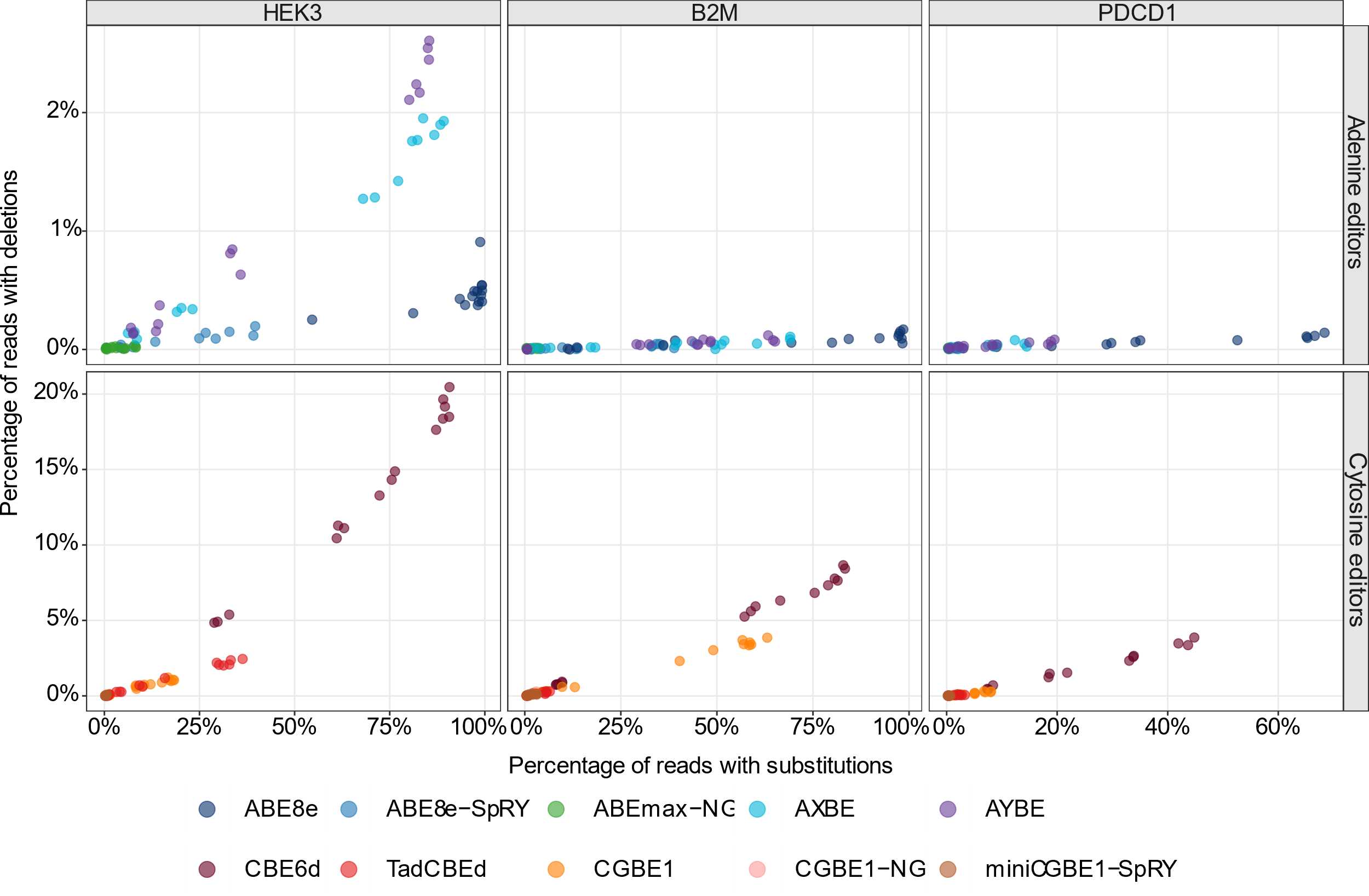

**Supplementary Figure 6: Relationship between substitution and deletion frequencies across dosages, editors, and target loci.** For each of the 10 base editors in the panel, the percentage of sequencing reads with a substitution (x-axes) is plotted against the percentage of reads with a deletion (y-axes), with each point representing one VLP dosage/replicate combination, for each of the editors (adenine editors (top panel) and cytosine base editors (bottom panel)) across the three target loci: HEK3, B2M, and PDCD1 (left – right subplots).

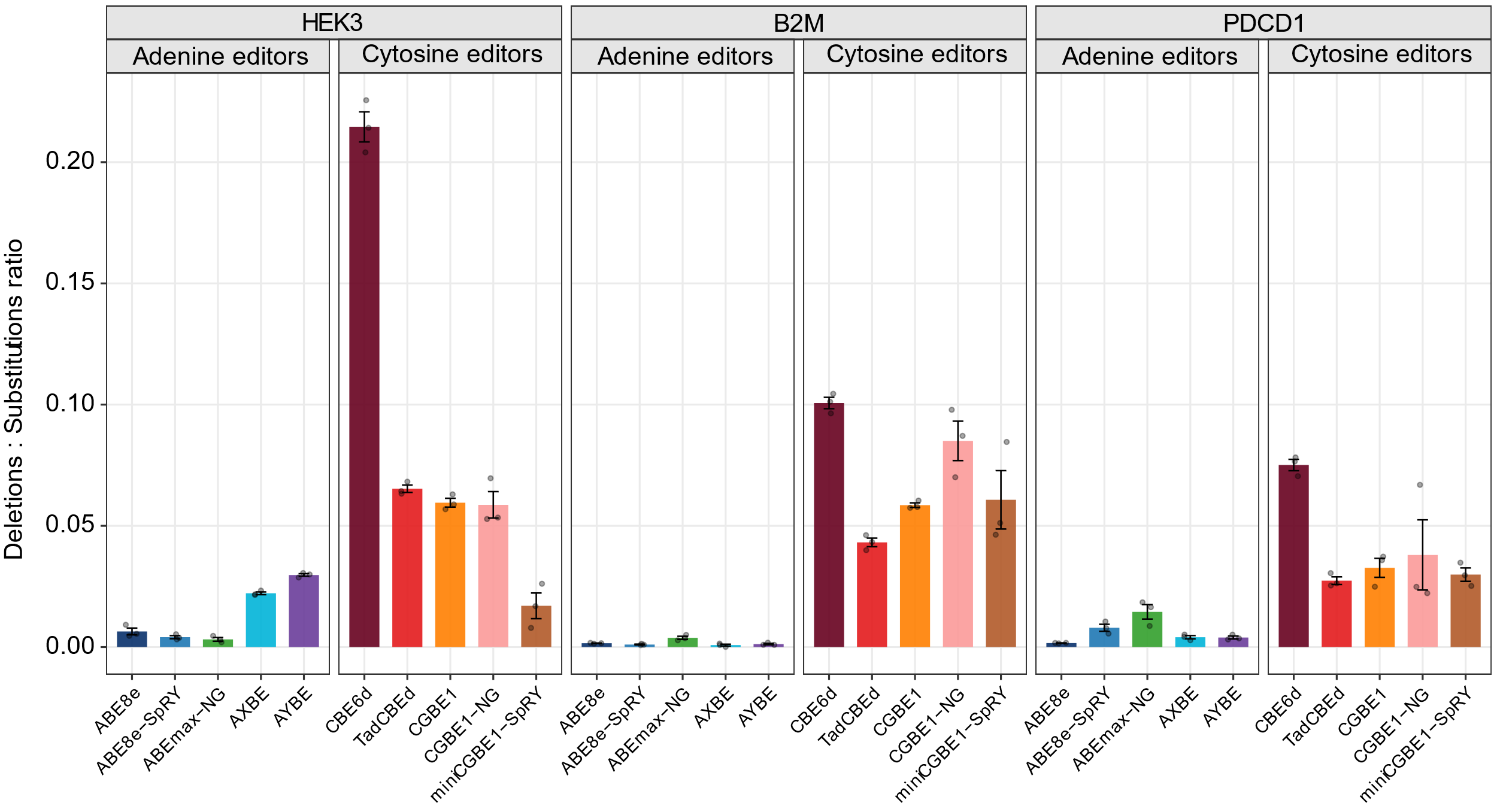

**Supplementary Figure 7: Deletion-to-substitution ratio across the base editor panel.** Ratio of deletion frequency to substitution frequency (y-axis) for each of the 10 benchmarked base editors (x-axis), for each target locus (columns). Bars represent the mean ratio across replicates; error bars denote SEM across replicates (n=3).

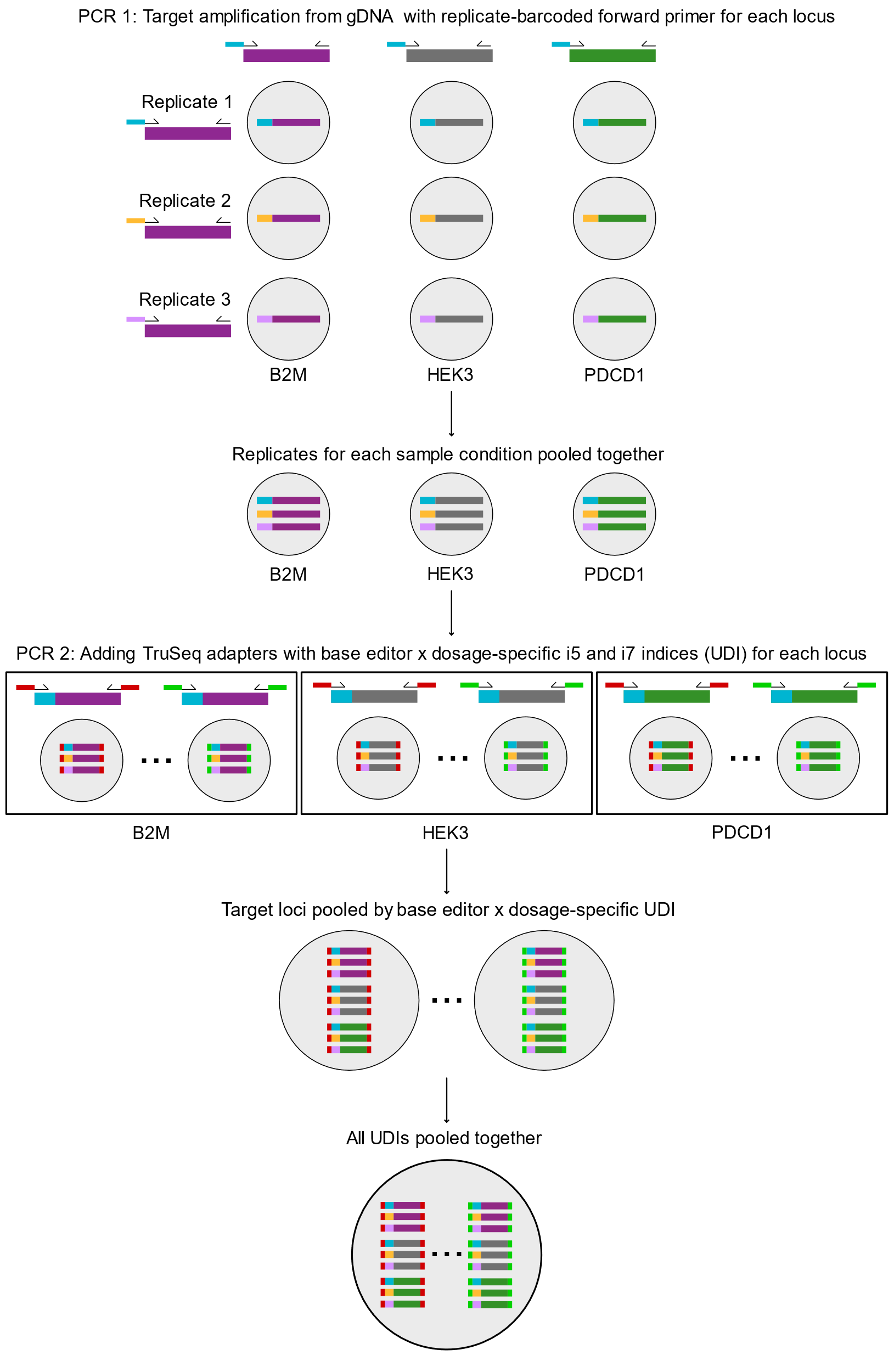

**Supplementary figure 8:** **Two-step, dual-barcoded PCR strategy for pooled amplicon library preparation.** Genomic DNA-containing cell lysate from each of the base editor × dosage condition for the three target loci (purple, gray, and green amplicons for B2M, HEK3, and PDCD1, respectively) and three biological replicates was amplified by PCR1 using replicate-specific forward primers (cyan, yellow, and pink barcodes for Replicates 1 to 3). Replicates were then pooled, and the locus-pooled amplicons underwent PCR2 to append TruSeq sequencing adapters together with a unique i5/i7 index pair assigned per sample condition (base editor × dosage; red and green primers, and others not shown). Finally, the indexed libraries from all three loci were combined into a single multiplexed sequencing pool. The reads are demultiplexed using Illumina indices to indicate base editor and dosage, the internal barcode indicating the replicate, and the locus indicating itself (see Methods).

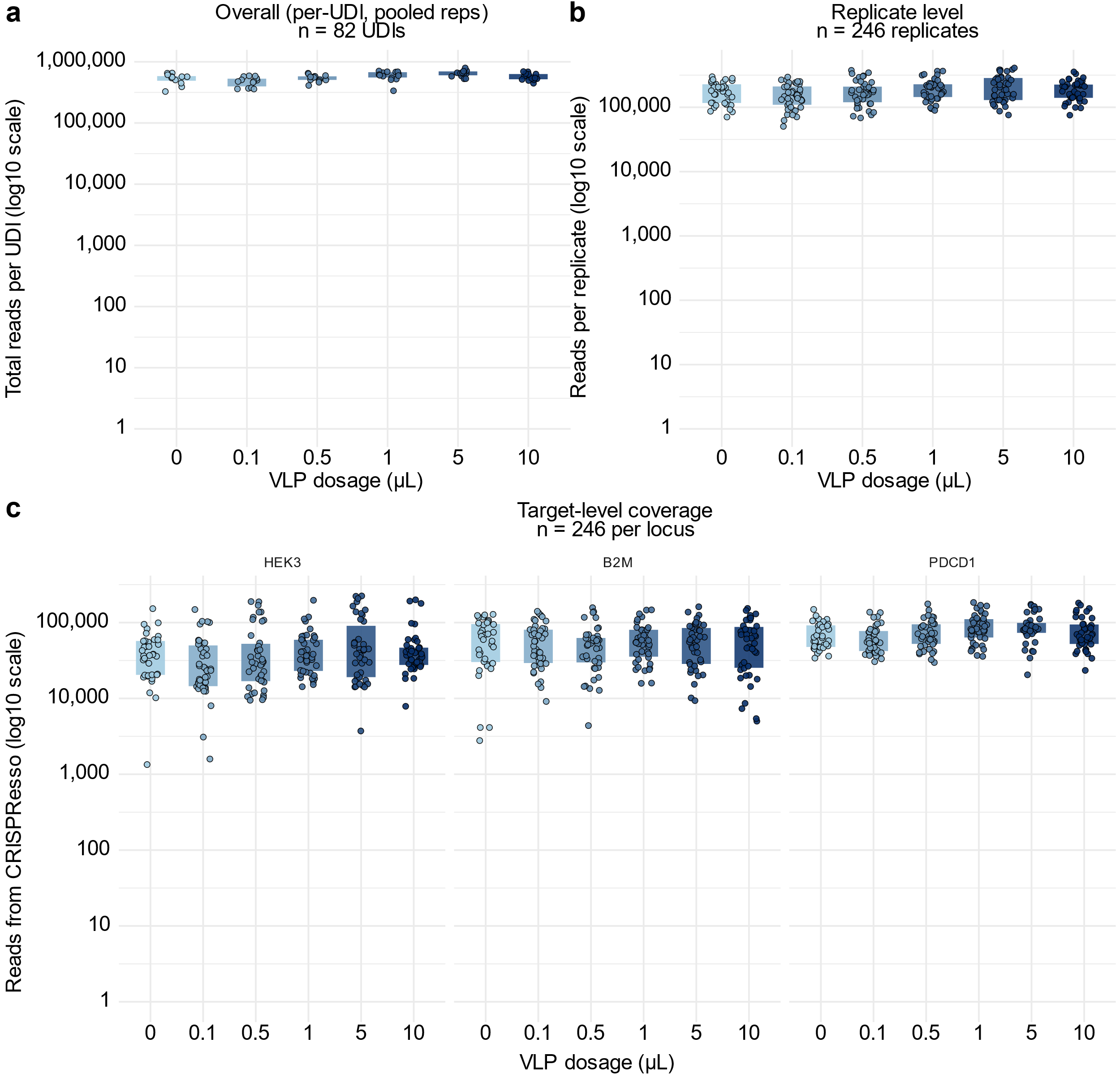

**Supplementary Figure 9: Sequencing read depth and coverage uniformity across VLP dosages, replicates, and target loci.** Boxplots showing the distribution of sequencing read counts (y-axes) at three levels of aggregation, colored by VLP dosage (x-axes) **(a)** Total reads per unique dual index (UDI), i.e., per sample condition (base editor × dosage) pooled across all three biological replicates and all three target loci (n = 82 UDIs). **(b)** Reads per condition prior to replicate demultiplexing (n = 246 replicate+base editor+dosage combinations; 82 UDIs × 3 replicates). **(c)** Target-level read coverage, by locus HEK3, B2M, and PDCD1 (left – right subplots; n = 246 replicate+base editor+dosage combinations per locus). 0 μL samples for AYBE and SPACE-ΔCBE were omitted from this graph as they did not have any sequencing reads.

| **Base editor** | **Target locus** | **Position** | **Total substitution (%)** | **C-to-T edits (%)** | **C-to-non-T edits (%)** |
| --- | --- | --- | --- | --- | --- |
| **CBE6d** | **HEK3** | **C3** | 33.41 | 87.2 | 12.8 |
|  |  | **C4** | 62.43 | 67.3 | 32.7 |
|  |  | **C5** | 37.39 | 64.0 | 36.0 |
|  |  | **C9** | 7.31 | 29.4 | 70.6 |
|  | **B2M** | **C2** | 1.51 | 51.6 | 48.4 |
|  |  | **C4** | 22.38 | 61.9 | 38.1 |
|  |  | **C6** | 43.29 | 59.2 | 40.8 |
|  |  | **C8** | 51.49 | 38.7 | 61.3 |
|  | **PDCD1** | **C4** | 3.41 | 50.1 | 49.9 |
|  |  | **C7** | 27.79 | 62.1 | 37.9 |
|  |  | **C8** | 5.48 | 36.0 | 64.0 |
| **TadCBEd** | **HEK3** | **C3** | 2.29 | 82.4 | 17.6 |
|  |  | **C4** | 9.95 | 59.4 | 40.6 |
|  |  | **C5** | 20.69 | 47.2 | 52.8 |
|  |  | **C9** | 0.51 | 16.4 | 83.6 |
|  | **B2M** | **C2** | 0.05 | 18.5 | 81.5 |
|  |  | **C4** | 1.15 | 44.5 | 55.5 |
|  |  | **C6** | 1.87 | 58.0 | 42.0 |
|  |  | **C8** | 2.71 | 26.5 | 73.5 |
|  | **PDCD1** | **C4** | 0.14 | 25.9 | 74.1 |
|  |  | **C7** | 1.33 | 57.7 | 42.3 |
|  |  | **C8** | 0.54 | 20.2 | 79.8 |

**Supplementary Table 1: Position-specific C-to-T and C-to-A/G editing frequencies for CBE6d and TadCBEd.** For each editable cytosine position within the protospacer at each of the three target loci (B2M, HEK3, PDCD1), total substitution frequency (percentage of sequencing reads with a substitution at that position) is reported alongside its breakdown into C-to-T edits and C-to-A/G edits, expressed as a percentage of total substitutions at that position. Only positions with detectable editing (≥1% total substitution frequency at least one editor) are shown.
